# Structured local mismatch placement and internal packaging in cardiomyocytes during hyperthermal sarcomeric oscillations

**DOI:** 10.64898/2026.03.26.714639

**Authors:** Seine A. Shintani

## Abstract

Neighboring sarcomeres during hyperthermal sarcomeric oscillations (HSOs) are not perfectly synchronous, yet their local nonuniformity is organized rather than random. Earlier reanalyses of the same continuous five-sarcomere recordings showed that local reconfiguration is dominated by minimal Hamming-1/one-link updates and that fast cycles can be aligned onto a shared rephasing compass whose clearest physiological translation is mismatch-pocket placement along the observed chain. Here we integrate those results with reduced-geometry analysis that asks how the same minimal update is internally distributed within the chain. Using seven living neonatal rat cardiomyocytes recorded at 500 frames/s, we converted each fast HSO cycle into one local coordination summary, extracted event-centered internal pre-to-post sarcomere-length change patterns after subtraction of the instantaneous five-sarcomere mean, and retained exact directed one-link transitions together with mismatch-pocket context. Continuous reduced-plane trajectories were geometrically complex, and total event displacement was not identical to redistribution span. Cross-cell alignment again revealed a common rephasing order, and aligned position most strongly predicted edge-biased mismatch placement. Reduced geometry added a distinct mesoscale layer: route classes formed an ordered redistribution-span axis from compact to broad internal redistribution, and even when exact one-link transition and pocket context were matched, events still separated into compact and extended redistribution profiles. Across common matched groups, the extended family showed broader span in 8 of 9 groups (median span difference +0.423; paired Wilcoxon *P* = 0.027). These findings support a structured-mismatch view of HSOs: the observed five-sarcomere chain reuses the same minimal local reconfiguration through more than one internal redistribution route.

**Significance statement:** Cardiac contraction must transform noisy local events into a stable beat. This study shows that local nonuniformity in living cardiomyocytes is structured at more than one mesoscale level. A shared rephasing compass organizes where a mismatch pocket tends to sit along an observed five-sarcomere chain, and reduced geometry shows that the same minimal local update can still be internally packaged through more than one redistribution route.

Graphical abstract.
An observed fast HSO window is condensed into cycle-wise local phase summaries. A topology-based circular coordinate and cross-cell alignment yield a shared rephasing compass. The clearest primary readout of that compass is mismatch-pocket placement along the observed five-sarcomere chain, whereas reduced geometry adds a secondary descriptive layer in which route packaging is ordered from compact to broad redistribution span.

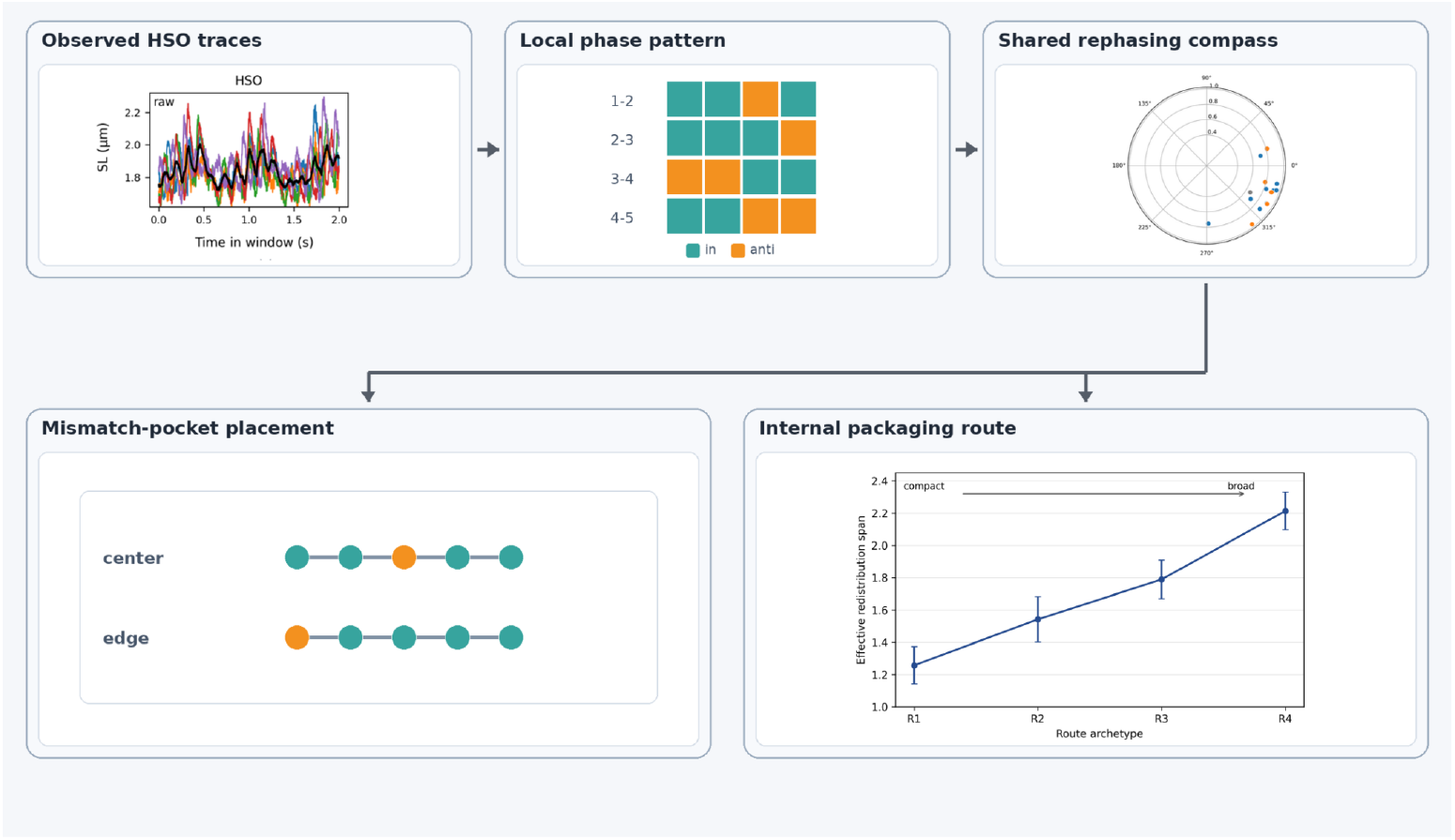

## 1 Introduction

Cardiac contractility is a multiscale problem. Molecular events are stochastic, yet the cell and the heart still generate an ordered beat. Reviews of cardiac contractility and myosin mechanics emphasize that this bridge must be understood across scales, from single molecules to whole hearts, with sarcomeres and myofibrils acting as key intermediate levels (Garg et al., 2024; Rassier and Månsson, 2025). Direct measurements also show that neighboring sarcomeres in living myocardium are heterogeneous in length and strain, that stretch can harmonize part of that heterogeneity, and that sarcomeric synchrony and asynchrony matter for ventricular pump function (Li et al., 2023; Lookin et al., 2023; Kobirumaki-Shimozawa et al., 2021, 2024). Local nonuniformity is therefore not just technical noise. The unresolved problem is how a cardiomyocyte preserves organized output while adjacent contractile elements are not perfectly identical.

Warmed cardiomyocytes provide a useful regime in which this issue becomes experimentally visible. Local heat pulses can trigger contraction without detectable calcium transients, microscopic heating can activate cardiac thin filaments directly, and thin-filament activation is strongly cooperative and mechanically regulated (Oyama et al., 2012; Ishii et al., 2019, 2020; Solís and Solaro, 2021; Park-Holohan et al., 2021). Hyperthermal sarcomeric oscillations (HSOs) are therefore not simply a failure of beating. They are a living-cell state in which a slower beat-scale envelope persists while rapid internal oscillation becomes easier to read (Shintani et al., 2015, 2020; Shintani, 2022, 2025).

Our earlier reanalysis of the same five-sarcomere recordings provided one anchor for this regime: local reconfiguration occupied a constrained 16-state neighboring-pair topology dominated by Hamming-1 transitions, and the segment-mean rapid signal was closely described by the product of local oscillation size and weighted synchrony, *A × R*_*w*_ (Shintani, 2026). A separate reanalysis of the same fast cycles showed that cell-wise circular coordinates can be aligned across cells into a shared rephasing order whose clearest physiological translation is mismatch-pocket placement along the observed chain. Those studies answered two different questions: what changes, and where mismatch is most readably positioned. What they did not explain was why cycles sharing the same coarse local context could still differ in their internal redistribution patterns. The present study addresses that remaining question by bringing reduced-geometry packaging into the shared-compass framework.

In plain language, we move from *what changes* and *where the mismatch pocket sits* to *how the same minimal update is internally distributed within the observed chain*. Reduced geometry is used here as a descriptive mesoscale packaging language rather than as a new output law. The central message remains simple. Local HSO nonuniformity is structured at more than one mesoscale level: a shared rephasing compass organizes where mismatch is positioned, and reduced geometry organizes how the same minimal reconfiguration is internally packaged within the five-sarcomere chain.

## 2 Results

For clarity, the main figures are ordered around five practical questions. Figure 1 defines the raw data and the common analytical unit. Figure 2 shows how a continuous reduced trajectory yields event-level local steps and why total event size is not the same as redistribution span. Figure 3 returns to the cycle-wise compass question and asks whether local states share a common order across cells and how that order is translated most readably. Figure 4 asks the reduced-geometry question: whether the same minimal one-link update is internally packaged in one way or in several. Figure 5 then ranks the resulting claims by robustness.

**Figure 1:**
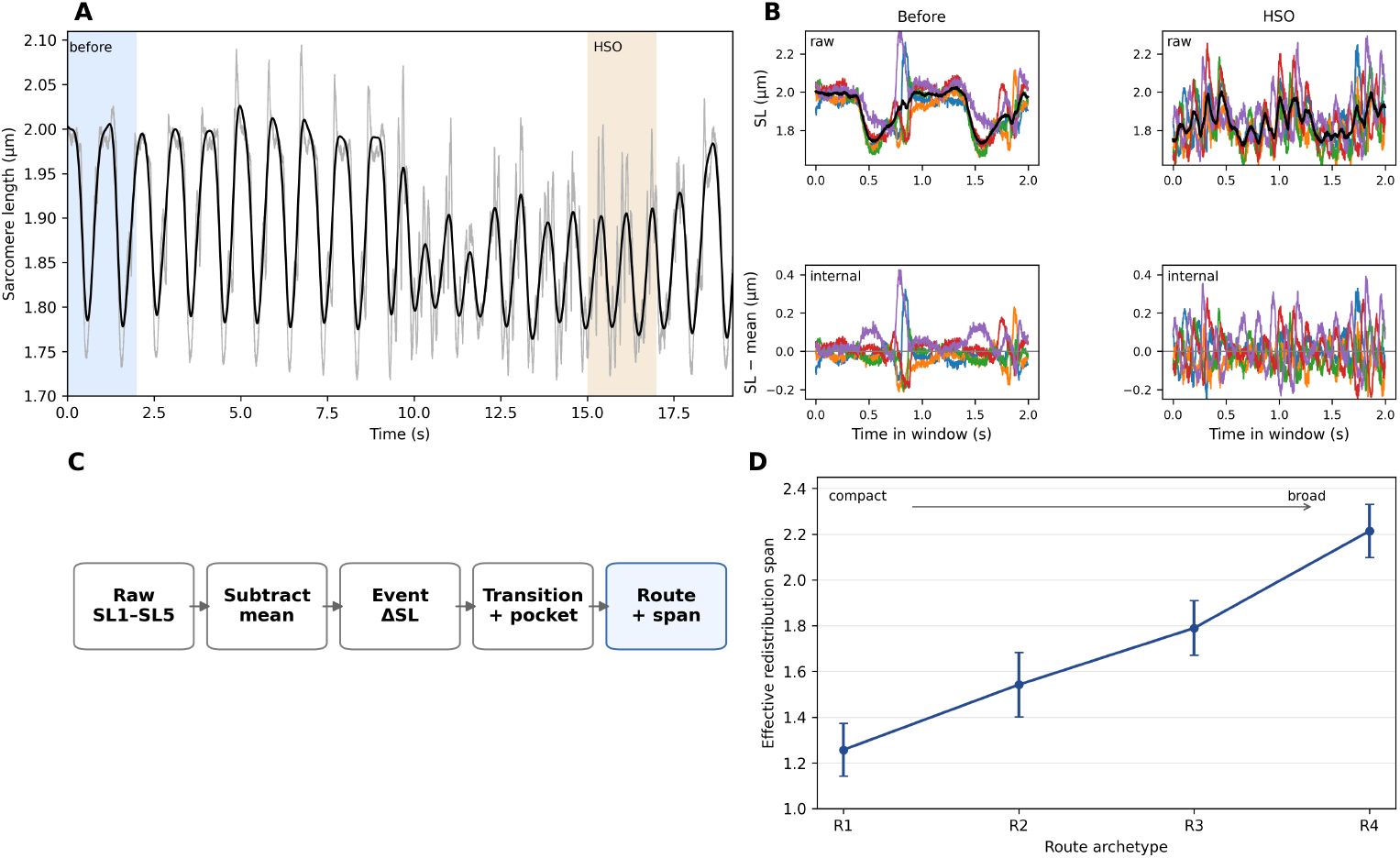
Raw five-sarcomere recordings and the analytical unit used in this study. Panel A shows a representative continuous recording with before-warming and HSO windows indicated. Panel B contrasts short before-warming and HSO windows in the raw traces and in the internal components after subtraction of the instantaneous five-sarcomere mean. Panel C summarizes the analysis ladder from raw traces to event-centered internal displacement, transition+pocket context, and route/span summaries. Panel D shows that route classes form an ordered redistribution-span axis from compact to broad internal redistribution.

**Figure 2:**
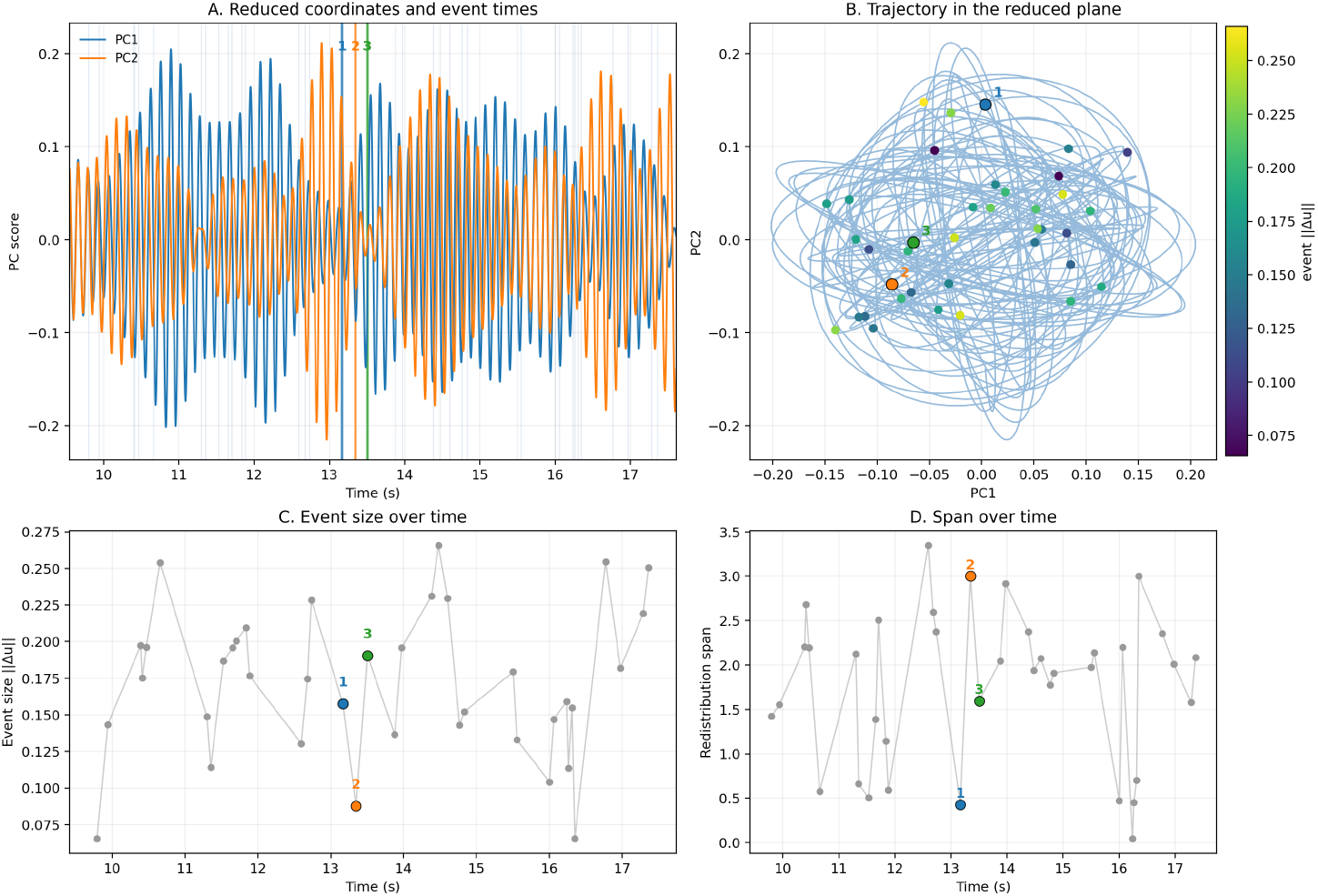
Continuous trajectory, event extraction, and the distinction between size and redistribution span. Representative reduced-plane trajectory from one HSO window. The time-domain panels show the reduced coordinates and event times, and the reduced-plane panel shows the same window as a continuous trajectory. Event-sized displacement and redistribution span are then plotted as separate summaries. The key message is that the continuous trajectory is geometrically complex, that local events can still be extracted from it, and that total event displacement is not identical to the breadth of the internal redistribution pattern.

**Figure 3:**
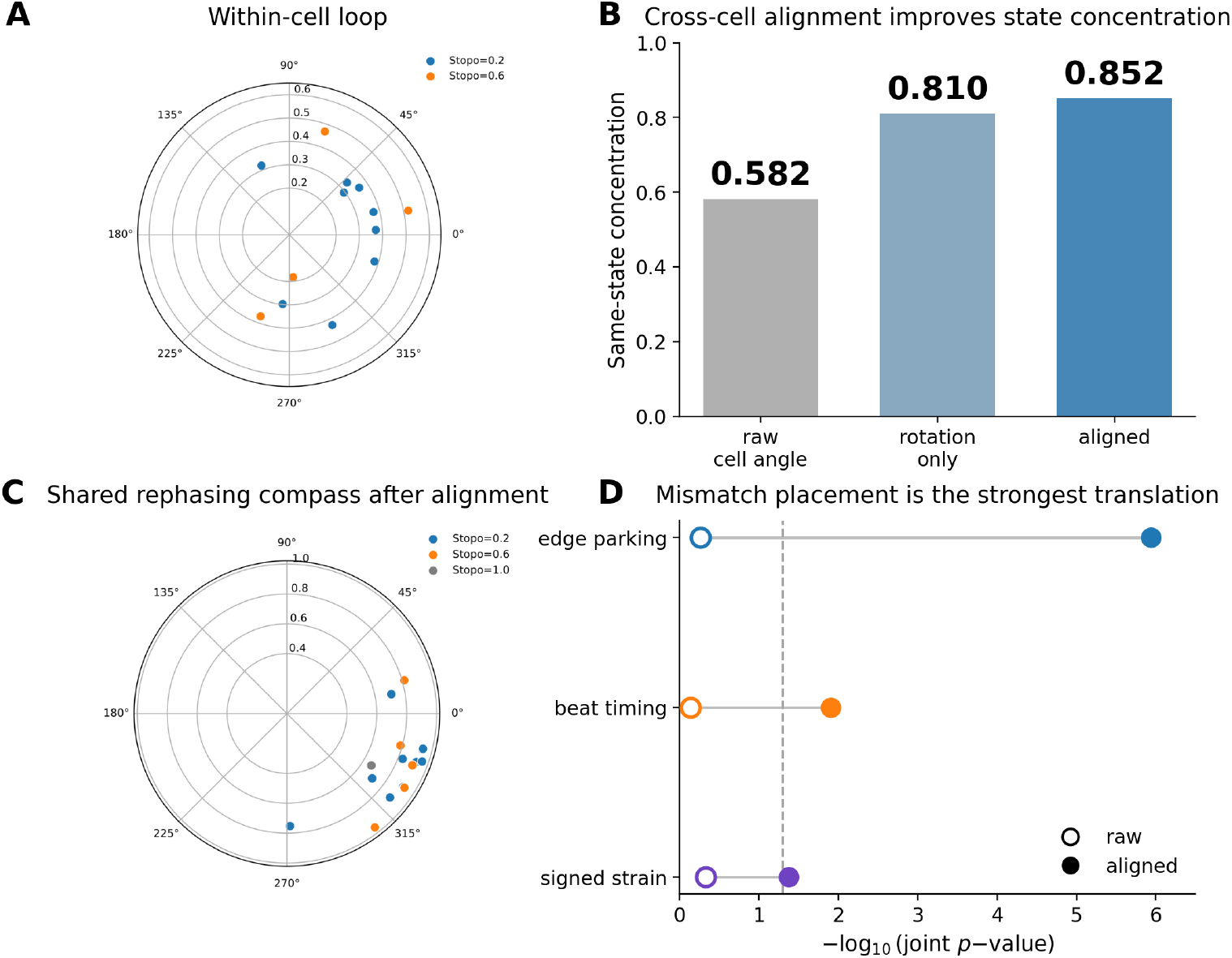
Within-cell order, cross-cell alignment, and mismatch placement. Panel A shows a within-cell circular-coordinate exemplar that underlies the pooled compass step. Panel B shows same-state concentration for raw cell-wise angles, a rotation-only control, and the fully aligned solution (0.582, 0.810, and 0.852, respectively). Panel C shows the pooled shared rephasing compass after alignment; it should be read as a common ordering rather than as an absolute physiological angle. Panel D compares candidate physiological translations before and after alignment. Mismatch placement / edge bias is the strongest and most robust translation, whereas beat timing and signed strain are weaker. Supplementary Figure S1 retains the broader within-cell prerequisite without duplicating this main-text pooled figure.

**Figure 4:**
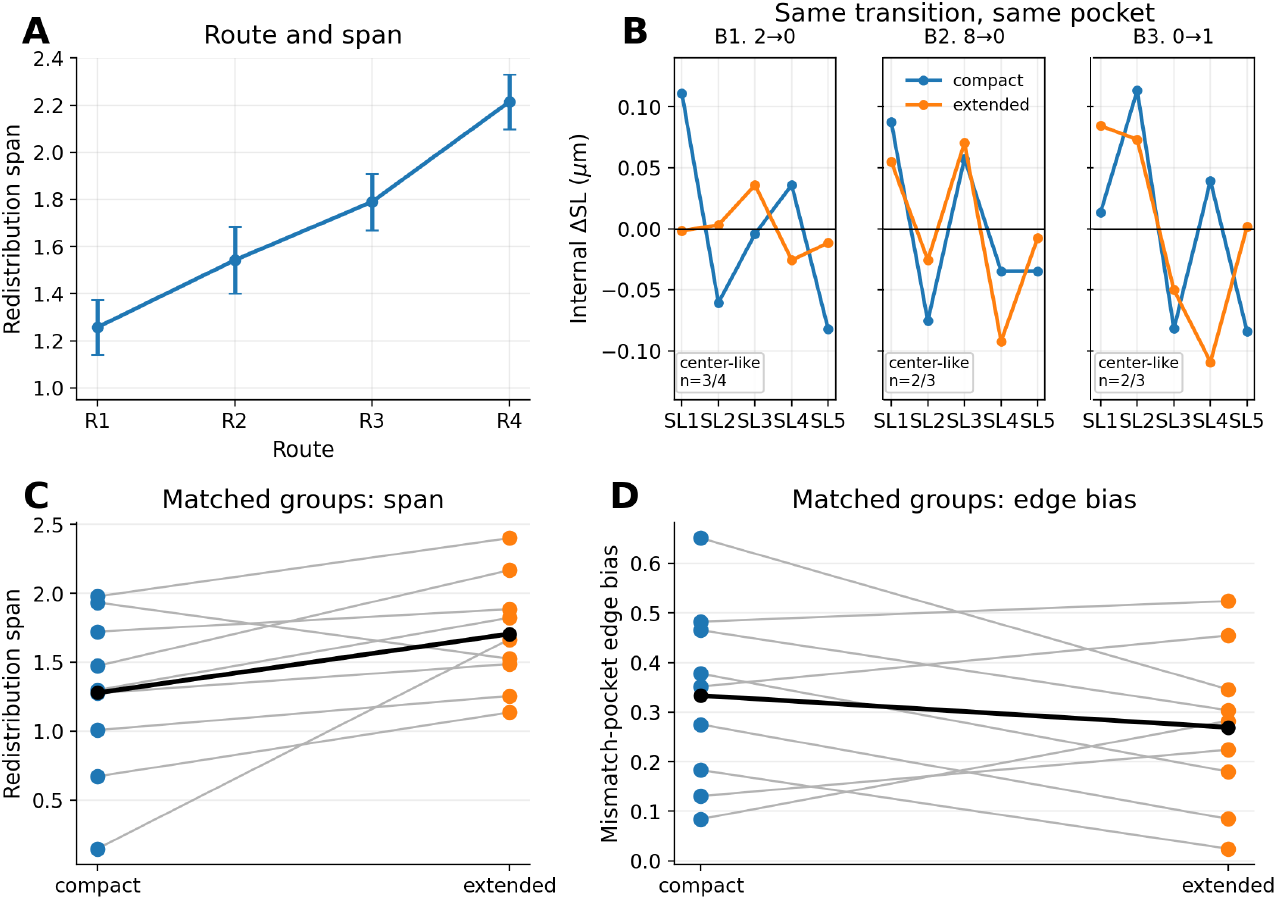
The same minimal update still uses multiple internal packaging routes. Panel A shows an ordered redistribution-span axis from compact to broad internal redistribution. Panel B shows representative matched examples in which compact and extended profiles coexist despite the same exact directed one-link transition and the same pocket context. Panels C and D summarize common matched groups: redistribution span shifts consistently toward the extended family, whereas mismatch-pocket edge bias does not. Reduced geometry is therefore used here as a mesoscale packaging descriptor rather than as a new output law.

**Figure 5:**
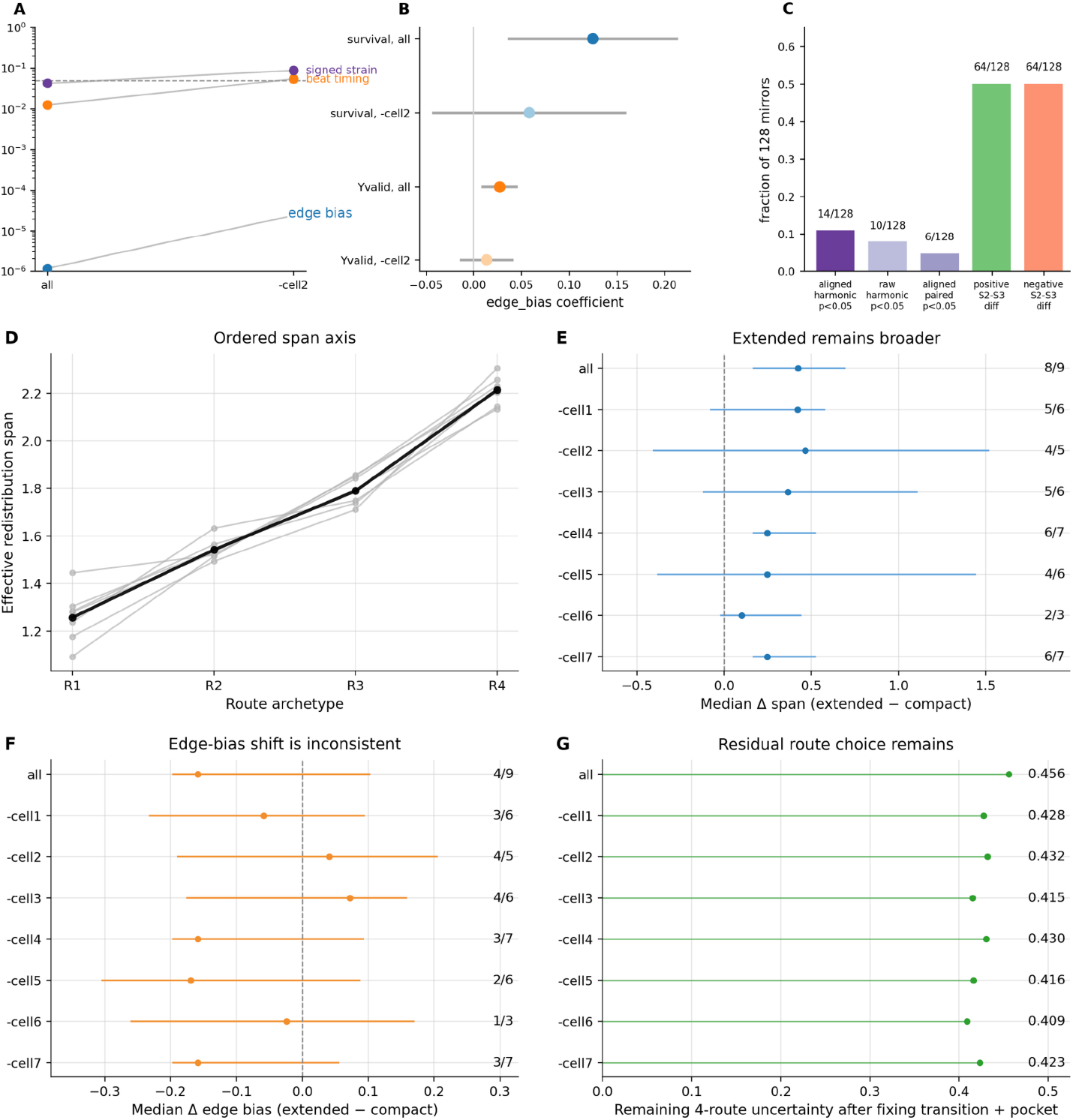
Robustness and claim hierarchy. Panels A–C rank the compass-side claims: mismatch placement is the strongest and most robust translation, whereas beat timing and signed strain are weaker. Panels D–G add leave-one-cell-out robustness for the route-packaging message. Across all seven leave-one-cell-out splits, the ordered route-span axis is preserved and residual route choice remains after strict transition-plus-pocket conditioning. Together, these panels justify the hierarchy used throughout the manuscript: mismatch placement is the strongest mesoscale readout, and route packaging adds a robust qualitative layer about how the same minimal update is internally distributed.

### 2.1 Raw data, analytical unit, and event-level packaging descriptor are defined from the same five-sarcomere recordings

Figure 1 defines the common analytical starting point for both the compass and packaging branches. The starting material is the same primary recording used in the earlier HSO papers: one continuous five-sarcomere line in each of seven cells, recorded at 500 frames/s. During HSOs, the slow beat-scale envelope remains visible, but faster internal oscillation becomes much more prominent. The before-warming window in Figure 1 is included only to orient the reader to the HSO regime; all shared-compass and route-packaging analyses below are restricted to fast HSO cycles.

Each fast HSO cycle is first converted into one local coordination summary. Four neighboring-pair relations define one dominant 16-state local pattern together with circular position, aligned circular position, pooled aligned sector, beat-phase position, and mismatch-placement summaries. This keeps every later analysis anchored to directly observed fast cycles rather than to a hidden-state model.

The packaging branch then adds one further step. We subtract the instantaneous five-sarcomere mean from the raw traces so that the analysis emphasizes redistribution within the observed chain rather than the slower common beat motion. From those mean-subtracted traces, event-centered internal pre-to-post sarcomere-length change patterns are extracted and summarized by reduced-geometry route and redistribution span. The analytical unit is therefore a directly observed minimal local update together with its measured internal redistribution pattern.

### 2.2 Continuous reduced-plane trajectories are complex, but local events can still be extracted and read by both size and redistribution span

This step is shown explicitly because route labels could otherwise look like an after-the-fact abstraction. Figure 2 therefore returns to the raw event stream and shows how the reduced trajectory is built from directly observed time series.

The reduced-plane trajectory is continuous and geometrically complex rather than a simple monotonic orbit. Event times can nonetheless be placed on that trajectory, making it possible to follow local steps within the larger HSO window. In the same time series, total event displacement and redistribution span are not identical. Large and small events are interleaved, and a relatively small total displacement can still be associated with a broader redistribution span. In biological terms, the size of a local step and the breadth of the compensating redistribution are different properties.

Figure 2 is therefore not presented as an orbit law. Instead, it explains why a packaging description is useful: the observed trajectory is too rich to summarize simply as small versus large motion, yet structured enough that event-centered local steps can be extracted and compared reproducibly.

### 2.3 Cross-cell alignment reveals a shared rephasing compass whose strongest physiological translation is mismatch placement

Before pooling across cells, the cycle-wise local states within each cell already occupied a loop-like order beneath the 16-state description. In the cycle-wise analysis of these same data, adjacent positions on that within-cell loop were enriched for Hamming-1 changes (0.504 versus 0.284 under a null ordering), and cycle-to-cycle angular drift was slower than expected by chance (mean |Δ*θ*| = 0.285 versus 0.844 rad under shuffle). We do not devote a separate main-text figure to the full within-cell loop detail, but its explanatory role is preserved here in the prose and in Supplementary Figure S1.

We next ask the pooled question. Because each cell has an arbitrary zero point and direction, raw circular coordinates cannot be pooled directly. We therefore aligned the cell-wise circles using shared local states as landmarks. After alignment, same-state concentration improved from 0.582 to 0.852 (*P* = 0.0013), whereas a rotation-only control improved concentration only partway to 0.810. In other words, the pooled structure is not already present in the raw cell-wise angles; it emerges when the circles are put onto a common compass. We therefore continue to use the term *shared rephasing compass*. As before, that aligned coordinate should not be read as an absolute physiological angle. It is a common ordering that makes local mismatch configurations comparable across cells.

We then asked what this common ordering means in ordinary physiological language. Among the candidate translations tested, mismatch placement along the observed five-sarcomere chain was the strongest. Aligned angle strongly predicted edge-biased mismatch placement (joint *P* = 1.15 × 10−6), whereas the raw-angle control was null (*P* = 0.55). Beat timing aligned more weakly, and signed strain was weaker and more fragile still. The accessible reading is therefore not simply that cycles differ by an abstract angle. The shared compass most clearly tells us where the local out-of-step pocket tends to sit within the chain.

### 2.4 The same minimal update still splits into multiple internal packaging routes

Having established that local HSO nonuniformity is organized by a shared compass and most readably expressed by mismatch placement, we then asked whether the same minimal update is internally realized in only one way or in several. Here, a one-link update means that only one of the four neighboring-pair relations changed between the before and after local states - the smallest possible local update along the observed five-sarcomere chain. Reduced geometry was introduced to answer what happens next inside the chain.

Route classes formed an ordered redistribution-span axis from compact to broader internal redistribution. In practical terms, R1 and R2 form a more compact/lower family, whereas R3 and R4 form a more extended family. The important point is not the label itself but what it summarizes: how broadly the compensating redistribution is spread within the observed chain.

The key test was deliberately strict. We fixed the exact directed Hamming-1 transition and the same mismatch-pocket context, and then asked whether events still separated into more than one internal redistribution pattern. The answer was yes. Representative matched groups show that compact and extended events can coexist under the same local transition and pocket context. Across common matched groups, the extended family showed broader redistribution span in 8 of 9 groups (median Δspan = +0.423, paired Wilcoxon *P* = 0.027), whereas edge bias itself did not shift consistently across those matched comparisons. The safest interpretation is therefore that route is not simply another name for edge-biased mismatch placement. Rather, route describes how the same minimal update is internally packaged after pocket placement has already been taken into account.

That strict matched-group result is consistent with a broader conditioning analysis. Total 4-route entropy across HSO single-link events was 1.987 bits. Conditioning on dominant state alone still left 86.1% of route uncertainty, conditioning on aligned sector alone left 96.3%, and conditioning on dominant state plus aligned sector together still left 66.0% (62.1% excluding cell 2). Common state+sector groups were therefore usually mixed rather than single-route classes (median entropy 1.750 bits). State and sector bias route usage, but they do not uniquely specify it.

### 2.5 Robustness analyses rank the claims and keep the hierarchy readable

Sensitivity analyses were used to rank the claims rather than to inflate them. On the compass side, mismatch placement remained the strongest and most robust translation, whereas beat timing weakened to a secondary result and signed strain weakened further. On the packaging side, leave-one-cell-out analyses preserved the qualitative ordering of route span and the general tendency for extended events to remain broader than compact events under the same matched-group rule. Across all seven leave-one-cell-out splits, the ordered span axis R1<R2<R3<R4 was preserved, and residual 4-route uncertainty after fixing exact transition plus pocket context remained substantial (0.409–0.432).

This ranking is important for how the manuscript should be read. The strongest shared-compass claim remains mismatch placement along the observed chain. The new reduced-geometry claim is that the same minimal update still has more than one internal packaging route and that this distinction is qualitatively robust. Beat timing, signed strain, and stricter exact-state formulations remain informative but are best treated as secondary or supportive.

## 3 Discussion

The central message of this study is that local HSO nonuniformity is structured at more than one mesoscale level. The earlier topology analysis established what the minimal update looks like: neighboring sarcomeres reconfigure mainly through Hamming-1 / one-link steps (Shintani, 2026). The compass analysis established where a local mismatch pocket tends to sit after cross-cell alignment. The present analysis adds a third layer: the same minimal local update is not internally packaged in a unique way.

That layered interpretation is biologically useful. It suggests that the observed five-sarcomere chain does not merely tolerate local mismatch as random noise. Instead, the chain appears to organize local mismatch in a structured way. One part of that structure is where the minority-phase pocket tends to be placed along the observed segment. Another is how broadly the compensating redistribution is spread when the same minimal update occurs. In plain language, the chain appears to have more than one way to redistribute or relay a local mismatch while the broader beat-scale rhythm persists.

This framework also helps clarify what the reduced-geometry branch is and is not. It is not introduced here as a new output law competing with *A×R*_*w*_. The amplitude–synchrony relation remains an explicit cycle-level description of the observed segment mean (Shintani, 2026). Nor is the aligned circular coordinate treated as an absolute physiological angle. It remains a common ordering. Reduced geometry instead provides a descriptive language for the internal packaging of the same minimal local update. That is why route packaging should be read as a mesoscale descriptor layered beneath the shared compass rather than as a replacement for it.

The physiological implication is a softer and more useful picture of local heterogeneity. Neighboring sarcomeres are neither perfectly synchronized nor hopelessly disordered. Instead, the data support structured local mismatch organization. This interpretation fits with a broader literature in which sarcomere heterogeneity is biologically meaningful rather than incidental, and in which local mechanics feed back onto activation and force generation (Li et al., 2023; Kobirumaki-Shimozawa et al., 2021, 2024; de Tombe et al., 2010; Campbell, 2011). The present work does not solve that entire multiscale bridge, but it identifies a measurable intermediate-scale organization within it.

The hierarchy of claims should remain explicit. The strongest result is still mismatch placement along the observed chain. Route packaging adds a new and qualitatively robust layer, namely that the same minimal reconfiguration can still be internally distributed in more than one way. Dominant state and aligned sector organize that route usage, but they do not uniquely specify it: even after both are fixed together, 66.0% of route uncertainty remains. Beat timing, signed strain, and stricter exact-state formulations are informative but do not yet carry the same level of confidence. Making this ranking visible is part of what keeps the manuscript readable and trustworthy.

Several limits define the appropriate level of interpretation. The present dataset is unusual in that it combines seven independent living-cell recordings, one continuous five-sarcomere segment per cell, and a frame rate of 500 frames/s. The main constraint is therefore not weak data quality but scope. The present work does not establish a whole-cell propagation law, nor does it identify the molecular origin of route choice. Longer line segments, independent datasets, and simultaneous measurements of local Ca2+, force, and titin-related mechanics (Tsukamoto et al., 2016) will be needed to test how general this mesoscale organization is and how it is produced.

In summary, HSOs in living cardiomyocytes reveal not only a constrained one-link topology and a shared mismatch-placement atlas, but also a remaining flexibility in how the same minimal reconfiguration is internally packaged within a five-sarcomere chain. That is the specific contribution of the reduced-geometry branch within the shared-compass framework.

## 4 Materials and methods

### 4.1 Study design, data source, and ethics

This study is a reanalysis of previously acquired sarcomere-length recordings from neonatal rat cardiomyocytes. No new animal experiments were performed for the present work. The original experimental procedures were approved by the Animal Experiment Committee of the Faculty of Science at the University of Tokyo and were conducted according to the institutional implementation manual, as reported previously (Shintani et al., 2014, 2015, 2020). The reanalyzed dataset consisted of seven cells, with one observed five-sarcomere line segment analyzed in each cell. One sarcomere trace (cell 2 sarcomere 2) had previously been flagged as a mean-length outlier in the earlier cycle-level quality-control review and was therefore monitored explicitly in sensitivity analyses rather than ignored. The shared-compass and reduced-geometry analyses were performed on fast HSO cycles; before-warming segments were used only for regime contrast and to orient the reader to the HSO state.

### 4.2 Original cell preparation and microscopic system

The original experimental procedures have been described in detail elsewhere (Shintani et al., 2014, 2015, 2020). Briefly, immature cardiomyocytes were isolated from 1-day-old Wistar rats, cultured under standard conditions, and imaged one day after *α*-actinin-AcGFP transfection. Measurements used an inverted fluorescence microscope with an oil-immersion objective, a specimen-plane pixel size of 150 nm, and a frame rate of 500 frames/s. A 1550-nm laser was used as a local heat source and a 488-nm laser was used to excite AcGFP-actinin fluorescence.

### 4.3 Signal decomposition, local-state definition, and cycle-wise local coordination summaries

For each cell, the HSO fundamental frequency was estimated from the HSO segment by spectral analysis of the root-mean-square internal signal. Each sarcomere trace was centered by subtracting the instantaneous local mean of the five observed sarcomeres and then band-pass filtered around the cell-specific HSO-centered band. Instantaneous phase and amplitude envelope were obtained from the Hilbert transform. Phase-valid intervals were defined from the mean high-frequency envelope using a hysteresis gate.

At each phase-valid time point, the five-sarcomere segment was represented by the four neighboring-pair phase relations 1–2, 2–3, 3–4, and 4–5. A neighboring pair was classified as in-phase when cos(Δ*ϕ*) ≥ 0 and anti-phase when cos(Δ*ϕ*) < 0. The four binary neighboring-pair relations yielded 16 possible local phase patterns. Fast HSO cycles were summarized as one row per cycle, storing the dominant local state together with circular coordinate, aligned circular coordinate, pooled aligned sector, beat-phase position, mismatch-placement bias, and local mechanical summaries.

### 4.4 Reduced-geometry event summaries, route labels, and redistribution span

To focus on internal redistribution, we subtracted the instantaneous five-sarcomere mean from the raw traces and worked in the resulting mean-zero internal coordinate system. Each HSO single-link event was then summarized by an event-centered internal pre-to-post sarcomere-length change pattern. The reduced-geometry basis decomposes the mean-zero five-sarcomere chain into four internal modes. For each event, squared-share composition across those four modes (share_sq1 to share_sq4) was used as a four-component summary of how strongly each internal mode contributed to the event.

We assigned each event to the route archetype carried most strongly by that event, namely R1 through R4 according to the mode with the largest squared share. This classification is intentionally descriptive and transparent. It asks which internal mode carries the event most strongly, not which cluster an unsupervised algorithm finds. We also summarized each event by an effective redistribution span, defined from the distance between the main shortening site and the center of the compensating positive block in the canonical internal pre-to-post pattern. This scalar is interpreted only as a descriptive summary of breadth.

### 4.5 Circular coordinate, cross-cell alignment, and translation variables

Within each cell, persistent-homology / circular-coordinate analysis was used to derive a loop-like local rephasing coordinate from the cycle-wise state cloud (de Silva et al., 2011). Because the raw circular coordinate has arbitrary origin and direction, cell-wise coordinates were aligned across cells using shared occupied local states as landmarks. For pooled summary plots only, the aligned circular coordinate was divided into four equal bins, denoted aligned sectors 1–4. These sectors were treated as summary bins rather than as additional biological states.

Mismatch-placement bias quantified how close the minority-phase pocket was to one end of the observed chain relative to its center. Beat-phase position located the same cycle on the slow-beat baseline. Signed end-to-end strain asymmetry summarized the difference between the two ends of the observed segment but was interpreted cautiously because it depends on left–right assignment.

### 4.6 Matched-group comparisons, robustness analyses, and statistics

To test whether route remained after local context was fixed, events were grouped by the same exact directed Hamming-1 transition and the same mismatch-pocket category. Representative matched groups were used for display, whereas common matched groups were summarized quantitatively when both route families were represented and the total number of events was at least five. Matched-group comparisons used paired Wilcoxon signed-rank tests. Ordered route trends and selected bridge analyses were summarized with cell-adjusted linear models.

Route uncertainty was summarized with entropy-based conditioning analyses, asking how much route uncertainty remained after conditioning on dominant state, aligned sector, their combination, pocket context, or exact transition. Sensitivity analyses were used to rank claims rather than to inflate them. We therefore repeated key summaries after excluding cell 2 and used leave-one-cell-out analyses to check whether the qualitative packaging message depended on any one cell.

## Data and code availability

The reanalyzed dataset, the figure-level summary tables used for the present analyses, and the custom analysis code used in this study are available from the corresponding author upon reasonable request.

## Declaration of generative AI and AI-assisted technologies in the writing process

During the preparation of this work, the author used ChatGPT (OpenAI) to improve the readability and language of the manuscript. After using this tool, the author reviewed and edited the content as needed and takes full responsibility for the content of the publication.

## References

Kenneth S. Campbell. Impact of myocyte strain on cardiac myofilament activation. Pflügers Archiv – European Journal of Physiology, 462(1):3–14, 2011. doi: 10.1007/s00424-011-0952-3.

Vin de Silva, Dmitriy Morozov, and Mikael Vejdemo-Johansson. Persistent cohomology and circular coordinates. Discrete & Computational Geometry, 45(4):737–759, 2011. doi: 10.1007/s00454-011-9344-x.

P. P. de Tombe, R. D. Mateja, K. Tachampa, Y. A. Mou, G. P. Farman, and T. C. Irving. Myofilament length dependent activation. Journal of Molecular and Cellular Cardiology, 48(5):851–858, 2010. doi: 10.1016/j.yjmcc.2009.12.017.

Ankit Garg, Kory J. Lavine, and Michael J. Greenberg. Assessing cardiac contractility from single molecules to whole hearts. JACC: Basic to Translational Science, 9(3):414–439, 2024. doi: 10.1016/j.jacbts.2023.07.013.

S. Ishii, K. Oyama, T. Arai, H. Itoh, S. A. Shintani, M. Suzuki, F. Kobirumaki-Shimozawa, T. Terui, N. Fukuda, and Shin’ichi Ishiwata. Microscopic heat pulses activate cardiac thin filaments. Journal of General Physiology, 151(6):860–869, 2019. doi: 10.1085/jgp.201812243.

S. Ishii, K. Oyama, S. A. Shintani, F. Kobirumaki-Shimozawa, Shin’ichi Ishiwata, and N. Fukuda. Thermal activation of thin filaments in striated muscle. Frontiers in Physiology, 11:278, 2020. doi: 10.3389/fphys.2020.00278.

F. Kobirumaki-Shimozawa, T. Shimozawa, K. Oyama, S. Baba, J. Li, T. Nakanishi, T. Terui, W. E. Louch, Shin’ichi Ishiwata, and N. Fukuda. Synchrony of sarcomeric movement regulates left ventricular pump function in the in vivo beating mouse heart. Journal of General Physiology, 153(11):e202012860, 2021. doi: 10.1085/jgp.202012860.

F. Kobirumaki-Shimozawa, K. Oyama, T. Nakanishi, Shin’ichi Ishiwata, and N. Fukuda. Asynchronous movement of sarcomeres in myocardium under living conditions: role of titin. Frontiers in Physiology, 15:1426545, 2024. doi: 10.3389/fphys.2024.1426545.

J. Li, J. Sundnes, Y. Hou, M. Laasmaa, M. Ruud, A. Unger, T. R. Kolstad, M. Frisk, P. A. Norseng, L. Yang, I. E. Setterberg, E. S. Alves, M. Kalakoutis, O. M. Sejersted, J.-T. Lanner, W. A. Linke, I. G. Lunde, P. P. de Tombe, and W. E. Louch. Stretch harmonizes sarcomere strain across the cardiomyocyte. Circulation Research, 133(3):255–270, 2023. doi: 10.1161/CIRCRESAHA.123.322588.

O. Lookin, P. de Tombe, N. Boulali, C. Gergely, T. Cloitre, and O. Cazorla. Cardiomyocyte sarcomere length variability: membrane fluorescence versus second harmonic generation myosin imaging. Journal of General Physiology, 155(4):e202213289, 2023. doi: 10.1085/jgp.202213289.

K. Oyama, A. Mizuno, S. A. Shintani, H. Itoh, T. Serizawa, N. Fukuda, M. Suzuki, and Shin’ichi Ishiwata. Microscopic heat pulses induce contraction of cardiomyocytes without calcium transients. Biochemical and Biophysical Research Communications, 417(1):607–612, 2012. doi: 10.1016/j.bbrc.2011.12.015.

S.-J. Park-Holohan, E. Brunello, T. Kampourakis, M. Rees, M. Irving, and L. Fusi. Stress-dependent activation of myosin in the heart requires thin filament activation and thick filament mechanosensing. Proceedings of the National Academy of Sciences of the United States of America, 118:e2023706118, 2021. doi: 10.1073/pnas.2023706118.

Dilson E. Rassier and Alf Månsson. Mechanisms of myosin II force generation: insights from novel experimental techniques and approaches. Physiological Reviews, 105(1):1–93, 2025. doi: 10.1152/physrev.00014.2023.

Seine A. Shintani. Hyperthermal sarcomeric oscillations generated in warmed cardiomyocytes control amplitudes with chaotic properties while keeping cycles constant. Biochemical and Biophysical Research Communications, 611:8–13, 2022. doi: 10.1016/j.bbrc.2022.04.055.

Seine A. Shintani. Chaordic homeodynamics: the periodic chaos phenomenon observed at the sarcomere level and its physiological significance. Biochemical and Biophysical Research Communications, 760:151712, 2025. doi: 10.1016/j.bbrc.2025.151712.

Seine A. Shintani. Constrained neighboring-sarcomere phase topology shapes mean HSO amplitude in living cardiomyocytes. bioRxiv, 2026. doi: 10.64898/2026.03.13.711515. Preprint.

Seine A. Shintani, K. Oyama, F. Kobirumaki-Shimozawa, T. Ohki, Shin’ichi Ishiwata, and N. Fukuda. Sarcomere length nanometry in rat neonatal cardiomyocytes expressed with α-actinin-AcGFP in Z discs. Journal of General Physiology, 143(4):513–524, 2014. doi: 10.1085/jgp.201311118.

Seine A. Shintani, K. Oyama, N. Fukuda, and Shin’ichi Ishiwata. High-frequency sarcomeric auto-oscillations induced by heating in living neonatal cardiomyocytes of the rat. Biochemical and Biophysical Research Communications, 457(2):165–170, 2015. doi: 10.1016/j.bbrc.2014.12.077.

Seine A. Shintani, T. Washio, and H. Higuchi. Mechanism of contraction rhythm homeostasis for hyperthermal sarcomeric oscillations of neonatal cardiomyocytes. Scientific Reports, 10(1):20468, 2020. doi: 10.1038/s41598-020-77443-x.

C. Solís and R. J. Solaro. Novel insights into sarcomere regulatory systems control of cardiac thin filament activation. Journal of General Physiology, 153:e202012777, 2021. doi: 10.1085/jgp.202012777.

S. Tsukamoto, T. Fujii, K. Oyama, S. A. Shintani, T. Shimozawa, F. Kobirumaki-Shimozawa, Shin’ichi Ishiwata, and N. Fukuda. Simultaneous imaging of local calcium and single sarcomere length in rat neonatal cardiomyocytes using yellow cameleon-nano140. Journal of General Physiology, 148(5):341–355, 2016. doi: 10.1085/jgp.201611604.

