## Supplementary Figures for "Structured local mismatch placement and internal packaging in cardiomyocytes during hyperthermal sarcomeric oscillations"

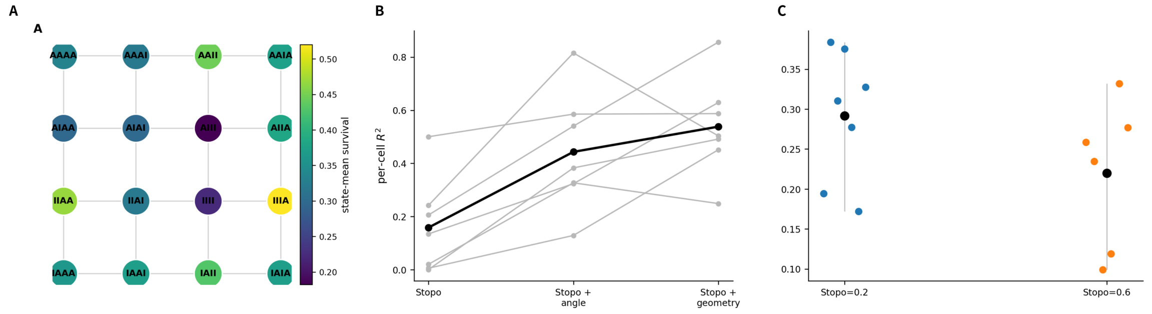


**Figure S1: Within-cell prerequisite for the shared compass without duplicating the main-text exemplar.** Panel A shows the Hamming-1 network of the 16 neighboring-pair patterns colored by within-cell loop order. Panel B shows that adding circular-angle or boundary-geometry terms improves the per-cell explanation of signal-survival ratio. Panel C shows that states sharing the same coarse synchrony score can still differ in within-cell survival range. This figure remains supplemental because it explains where the shared compass comes from, whereas the main text uses Fig. 3 only for the pooled cross-cell step.

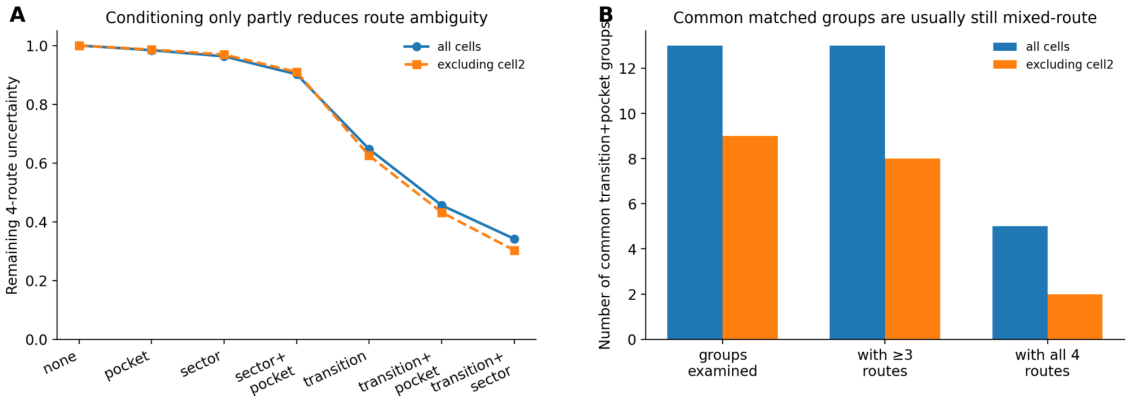


**Figure S2: Route ambiguity is reduced but not eliminated by coarse conditioning.** (A) Remaining 4-route uncertainty after conditioning on no label, pocket context, aligned sector, sector+pocket, exact transition, transition+pocket, or transition+sector. Conditioning reduces uncertainty, but substantial route choice remains even under the stricter transition-based summaries shown here. The main text separately reports the complementary dominant-state-plus-aligned-sector result from the same dataset. (B) Common matched groups are usually still mixed-route rather than single-route classes, even after exact transition and pocket category are fixed together.

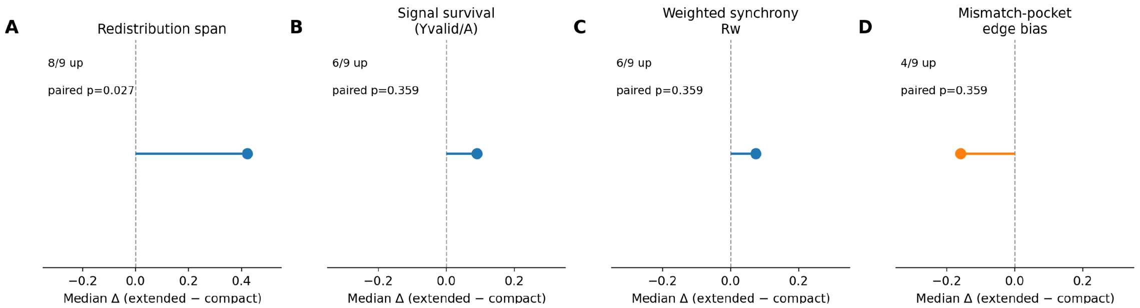


**Figure S3: Headline matched-group summaries support redistribution span as the clearest separator.** Paired summaries across common matched groups sharing the same exact transition and the same pocket context. Redistribution span shows the clearest compact-versus-extended separation (8/9 groups upward; paired Wilcoxon $P = 0.027$), whereas signal survival, weighted synchrony, and mismatch-pocket edge bias shift less consistently. This figure is kept supplemental because it supports the main span claim without duplicating the representative matched examples already shown in Fig. 4.

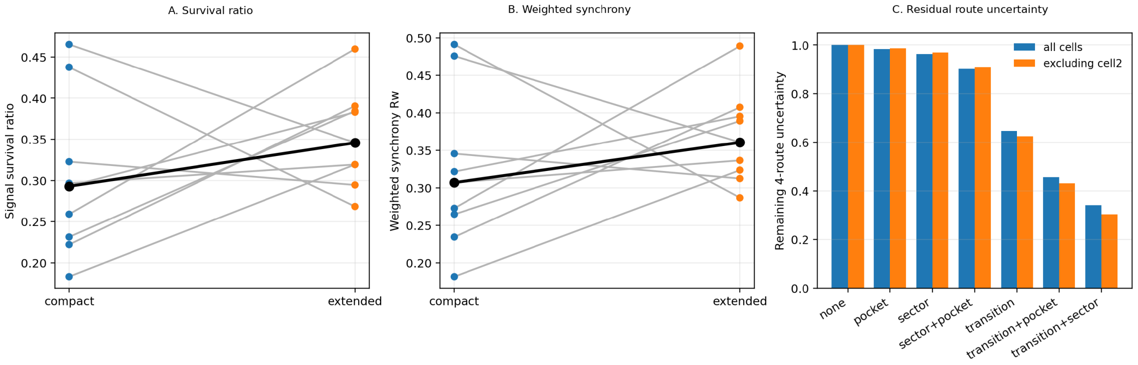


**Figure S4: Supportive raw paired values and conditioning series.** Panels A and B show the raw compact-versus-extended paired values for signal survival and weighted synchrony across the common matched groups. Panel C shows the full conditioning series for residual route uncertainty in all cells and after excluding cell 2. This figure remains supplemental because it provides supportive detail rather than a new headline claim.
